# PySeq2500: An open source toolkit for repurposing HiSeq 2500 sequencing systems as versatile fluidics and imaging platforms

**DOI:** 10.1101/2021.08.27.457864

**Authors:** Kunal Pandit, Joana Petrescu, Miguel Cuevas, William Stephenson, Peter Smibert, Hemali Phatnani, Silas Maniatis

## Abstract

Fluorescence microscopy is a key method in the life sciences. State of the art -omics methods combine fluorescence microscopy with complex protocols to visualize tens to thousands of features in each of millions of pixels across samples. These -omics methods require precise control of temperature, reagent application, and image acquisition parameters during iterative chemistry and imaging cycles conducted over the course of days or weeks. Automated execution of such methods enables robust and reproducible data generation. However, few commercial solutions exist for temperature controlled, fluidics coupled fluorescence imaging, and implementation of bespoke instrumentation requires specialized engineering expertise. Here we present PySeq2500, an open source Python code base and flow cell design that converts the Illumina HiSeq 2500 instrument into an open platform for programmable applications. Customizable PySeq2500 protocols enable experimental designs involving simultaneous 4-channel image acquisition, temperature control, reagent exchange, stable positioning, and sample integrity over extended experiments. To demonstrate accessible automation of complex, multi-day workflows, we use the PySeq2500 system for unattended execution of iterative indirect immunofluorescence imaging (4i). Our automated 4i method uses off-the-shelf antibodies over multiple cycles of staining, imaging, and antibody elution to build highly multiplexed maps of cell types and pathological features in mouse and postmortem human spinal cord sections. As demonstrated here, PySeq2500 enables non-specialists to develop and implement state of the art fluidics coupled imaging methods in a widely available benchtop system.

## Introduction

Imaging of fluorescently labeled biological materials underpins a wide range of state of the art techniques^1–7^. Many of these methods generate data over multiple cycles of chemistry and imaging. Robust and reproducible execution of these methods requires mechanical stability of the sample as well as precise and repeatable stage movements, temperature control, and reagent exchange. Due to a lack of commercially available instrumentation that meets these needs, researchers typically develop bespoke solutions for their specific applications. The engineering, programming, and optics expertise required to develop such bespoke solutions as well the high cost of acquiring and maintaining these systems present a substantial barrier to entry for many advanced fluorescence imaging applications. Moreover, the resulting non-uniformity of instrumentation across labs presents difficulties for replication and comparative analysis of data. Therefore, in order to enable the broader life science research community to adopt state of the art -omics methods based on fluorescence microscopy, an accessible open source toolkit of software, hardware, and protocols is desirable. We sought to address this need using hardware that is already available at many research institutions and that can be adapted for a variety of automated reagent handling and fluorescence imaging applications with minimal modification.

Illumina HiSeq instruments were once the workhorses of DNA sequencing, with over 2000 instruments in use as of 2018 (Illumina Investor Presentation, May 3, 2018). However, following the release of the NovaSeq 6000 instrument, Illumina announced it will no longer support HiSeq systems after February 2023. As a result, it is now possible to purchase used HiSeq 2500 systems on the secondary market for a fraction of their original price, making them appealing to be repurposed for novel applications. Formerly discontinued Illumina sequencing platforms have already been repurposed: the launch of the HiSeq family of instruments was accompanied by obsolescence of Illumina’s previous flagship instrument, the Genome Analyzer II (GAII), which became available on the secondary market and was adapted for custom applications^8–14^. Illumina’s currently available MiSeq system has also been modified for biological applications beyond sequencing^7,15^, though the modifications were less extensive than those implemented on the GAII by the Greenleaf group. The GAII was an open system with editable sequencing recipes and reagents that could be swapped out^16^. Internally, the GAII hardware consisted largely of off-the-shelf components. These factors made the GAII well suited for modification and custom applications by non-specialist research groups. In contrast, the HiSeq instrument family incorporates custom hardware, hard-wired control software, and barcoded reagents, making it less easily adapted for alternative applications^17^. However, given the imminent end of support for the HiSeq line, an easily implemented method for repurposing these instruments would make them powerful tools for automating a wide range of assays.

We present PySeq2500, an open source software toolkit for repurposing HiSeq 2500 systems to perform automated reagent exchange and fluorescence based imaging protocols. PySeq2500 controls and synchronizes all components within the HiSeq 2500 system, including the x, y, and z stages, microscope objective, pumps, valves, cameras, optics, and stage temperature control. An editable configuration file allows modification of HiSeq component settings and a customizable protocol file specifies the sequence of liquid handling, temperature, and imaging steps to be conducted. We also provide a novel sample flow cell design that is assembled from inexpensive commercially available materials. This flow cell integrates with the fluidics of the HiSeq 2500 and is compatible with epifluorescence imaging. The only hardware modifications required to adapt the HiSeq 2500 is the removal of 3 Phillips head screws to accommodate the custom flow cell.

To demonstrate the use of PySeq2500 for automated chemistry and imaging, we have developed an automated protocol for hi-plex staining and imaging of intact tissue sections using iterative indirect immunofluorescence imaging (4i). Unlike other spatial proteomics methods^5,18,19^, 4i enables highly multiplexed protein imaging in cultured cells and tissues using widely available conventional antibodies^4,20^. 4i uses a radical scavenging buffer during imaging to prevent photo-crosslinking of primary antibodies to their target antigen. This allows for gentle antibody elution in a mildly denaturing buffer, preserving sample integrity ^4,20^. Through iterative cycles of staining, imaging, and antibody elution, 4i can map up to dozens of proteins in a single sample^4^. We have automated this process using PySeq2500 to maximize the sample throughput, the number of proteins visualized, and the reproducibility of protein expression patterns across tissue sections. PySeq2500 automation enables unattended execution of multi-day, multi-cycle 4i experiments.

To highlight the versatility and robustness of our modified 4i protocol and flow cell, we used PySeq2500 to conduct automated 4i imaging of human and mouse spinal cord tissue sections. Our custom flow cell design provides a large surface area, accommodating multiple cm^2^ scale tissue sections on a single flow cell. Additionally, two flow cells can be processed in parallel, with chemistry performed on one flow cell while the second is being imaged. This parallel processing enables experimental designs that minimize batch effects between samples. We demonstrate the repeatability of our PySeq2500 automated 4i protocol by comparing staining of spinal cord sections from the SOD1-G93A mouse model of amyotrophic lateral sclerosis (ALS) and nontransgenic control animals processed in parallel. We also demonstrate the use of our automated 4i approach to visualize pathological features and cell types in postmortem spinal cord samples from subjects with ALS.

Through these experiments, we have characterized the performance of all HiSeq 2500 components. We provide an easy to use Python toolkit to control individual HiSeq2500 components through customized recipes that can be adapted for a variety of applications requiring robust automation.

## Results & Discussion

### HiSeq 2500 components

The Illumina HiSeq 2500 is a 4 color widefield optical epifluorescence microscope with integrated 3-axis motorized stages, temperature control, and liquid handling that can assay two flow cells in parallel. PySeq2500 flow cells are aligned to fluidic ports integrated within the stage and are held in place by vacuum. Reagents are drawn through flow cells by dedicated 8 barrel syringe pumps (250 uL per barrel). The on board refrigerator is capable of storing and sipping from up to 18 reagents per flow cell. A 24 port selector valve enables switching between reagents (ports 9, and 20-24 are unused in sequencing applications but are used by PySeq2500 to maximize reagent ports) (Figure 1A). Chemistry can be performed on both flow cells in parallel since the liquid handling system is duplicated for each flow cell. Flow cells are held to the stage with vacuum. The stage is moved in the x direction by a stepper motor with sub micron precision and in the y direction by a linear servo motor with a non-contact high precision linear encoder that provides 10 nm resolution. The stage can be tilted and moved in the z direction by three stepper motors. Samples are imaged using a 0.75 NA 20X air objective. The objective can be moved in the z direction by a piezoelectric linear actuator with 0.2 nm resolution. Samples within flow cells are excited by 2 liquid cooled lasers, with laser power modulated by the laser control hardware or by optical density filters (532 nm line up to 2W and 660 nm line up to 1W). Emitted light from the samples is filtered (558-32 nm, 610-60 nm, 687-20 nm, and 740-60 nm) (Figure 1B) and directed to 4 separate liquid cooled Time Delay Integration (TDI) line scanning CCD cameras. Each camera consists of 12 μm × 12 μm pixels with a field of view (FOV) of 769 μm x 6 μm. Temperatures of flow cells can be independently controlled by a combined peltier (ΔTmax 50°C) and liquid cooling system within the stage. To demonstrate experimental temperature control, A and B flow cell stage temperatures were changed to various setpoints from 20 to 55 °C using PySeq2500 and measured with an external thermocouple (Figure 1C).

**Figure 1.**
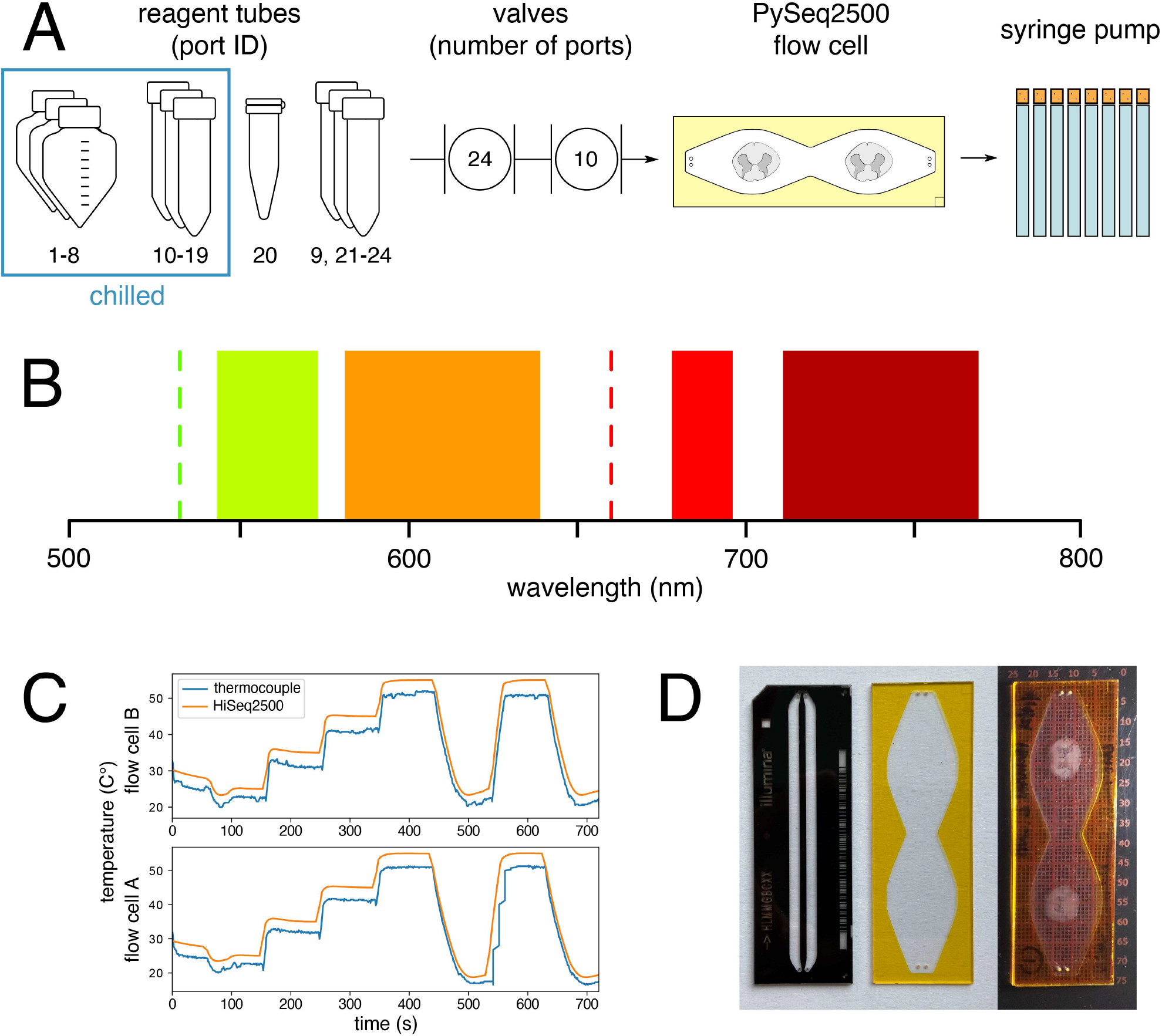
PySeq2500 control of HiSeq 2500 components. (A) The fluidic path is duplicated and independently operated for each flow cell (A and B). Reagents are drawn from their respective tubes into each flow cell by an 8 × 250 ***µ***L barrel syringe pump. Reagents are selected using a 24 port rotary valve with only 19 ports fully plumbed on a stock HiSeq 2500. With minor modifications, reagent lines can be added to ports 9 and 21-24. Up to 18 reagents per flow cell can be stored on the onboard chiller. A 2 hole stage inlet or 8 hole stage inlet is selected with the 10 port rotary valve. (B) Fluorescent markers are excited with lasers at 532 nm and 660 nm and emitted light is detected simultaneously at 558-32 nm, 610-60 nm, 687-20 nm, and 740-60 nm with bandpass filters. (C) A and B flow cell stage temperatures can be independently heated and cooled to various setpoints (orange) as measured using an external thermocouple (blue). (D) From left to right, an Illumina HiSeq 2500 8 lane flow cell, an empty PySeq2500 flow cell, and human spinal cord sections enclosed in a PySeq2500 flow cell atop a slide ruler used to measure ROIs to be imaged.

### A novel flow cell for automated reagent exchange and imaging

We designed a novel sample flow cell using inexpensive, accessible materials that is compatible with the fluidics of the HiSeq 2500. Given the upright configuration of the HiSeq imaging components, the PySeq2500 flow cell minimizes the distance between the sample and the objective, while ensuring a uniform headspace between the sample and the coverglass for reagent flow. It can also withstand a wide range of temperatures and is physically robust over multiple iterative cycles of chemistry and imaging. The flow cell is built around a standard glass microscope slide drilled with holes that align with the fluidics ports embedded in the HiSeq stage (Supplementary Figure 1A). The flow cell chamber is cut out of polyimide tape with double sided adhesive backing (Supplementary Figure 1B) and adhered to the glass slide with the outer protective film intact. After tissue cryosectioning and postfixation, flow cell assembly is completed by removing the outer protective film and adhering a coverglass (Figure 1D).

### PySeq2500 automation of reagent handling

The PySeq2500 software enables both interactive and automated control of the HiSeq 2500. Python modules control the individual components within the instrument to perform component specific tasks, such as checking the position of each stage. Higher level Python modules synchronize component level actions to perform complex system tasks, such as imaging a region of interest (ROI). The instrument can be controlled interactively by commands entered within a Python interpreter. Alternatively, a configuration file can be input to the software to automate execution of a specified protocol and to output log and image files. The configuration file specifies the protocol, valve port addresses of reagents, imaging ROIs, and additional settings of the HiSeq instrument.

Protocols are written as a sequence of 6 possible commands: PORT, PUMP, TEMP, IMAG, HOLD, and WAIT. The PORT command moves the 24 port selector valve to the specified reagent. The PUMP command draws in the specified volume in µL at a flow rate specified in the configuration file. The TEMP command changes the stage temperature of the flow cell to the specified temperature in °C. The IMAG command images ROIs on the flow cell as defined in the configuration file at a specified number of z planes. The HOLD command stops further actions on the flow cell until the specified time in minutes has elapsed. The WAIT command stops further action on the flow cell until the second flow cell starts a specified command, allowing coordination of imaging between flow cells.

### PySeq2500 automation of imaging

The HiSeq 2500 system uses TDI CCD cameras, which maximize the image signal and minimize noise by integrating signal from the same sample region over the time it takes for that region to traverse the TDI array, effectively extending exposure time. Given that the pixels of the CCDs in the HiSeq are laid out in 128 lines and the signal is integrated across them, noise arising from the readout process scales by a factor of 1 while the signal scales by a factor of 128. This detector configuration maximizes the signal to noise ratio but makes it difficult to obtain an interpretable live image since the FOV is a line. Therefore, instead of scanning the stage to locate tissue sections, the positions of samples on the flow cell are measured off the instrument and defined in the configuration file as ROIs.

Three dimensional ROIs within flow cells are imaged by first positioning the stage so that the corner of the ROI is just outside the FOV of the objective. The camera triggers are then armed and the y stage is moved at a constant velocity to scan the entire length of the sample. As the y stage moves, triggers are sent to the cameras which integrate the signal over space to form high signal to noise 12 bit images. After the scan is complete, the y stage is returned to the original position and the routine can be repeated at incremental objective z positions to build a volumetric tile. Once a tile is completed, the routine is repeated at incremental x positions to image the full volume/area of a ROI. Imaging ROIs of 3 × 3 mm and 8.5 × 10.5 mm with 10 focal planes was accomplished in approximately 40 and 155 minutes respectively.

The initial objective position in a z stack is calculated from the optimal objective position of the ROI, obtained with one of the following autofocus routines: full, full once, partial, partial once. The partial routines perform the autofocusing procedure on a single tile at the center of the ROI in the x direction while the full routines use the entire ROI to calculate the optimal objective position. The partial procedures should be used on ROIs larger than 5 × 5 mm to minimize focusing time given the high probability that the center tile of the ROI will overlap with the sample, while the full routines should be used on ROIs smaller than 5 × 5mm. The full once and partial once routines reuse the optimal objective position calculated in the first round for subsequent imaging cycles to minimize focusing time and preserve the correspondence of objective positions across imaging cycles. If the stage is moved in the z direction between imaging rounds, the full or partial routines, which recalculate the optimal objective position before each imaging cycle, should be used. Alternatively, a manual focus routine can be specified where an objective stack from the central FOV of the ROI is displayed to the user and the experiment is paused until the user inputs the frame that they judge to be most in focus.

### Validating automated 4i protocol using PySeq2500

To determine the optimal number of elution buffer incubations to fully remove bound primary and secondary antibodies from fresh frozen tissue sections in our automated 4i protocol, we conducted an elution series experiment. We first performed automated staining and imaging of tissue sections. We imaged the stained tissue sections after each of 6 incubations with 4i elution buffer^4^. After the first elution cycle, there is a dramatic decrease in GFAP and ELAVL2 signal as measured by pixel intensity. By cycle 3, the antibody signal is indistinguishable from the tissue background signal (Supplementary Figure 2). To ensure complete elution across all antibodies and tissue types, we used 4 elution incubation steps for mouse spinal cord sections and 5 elution incubation steps for postmortem human spinal cord sections in subsequent experiments.

To characterize the stability of data generated using our automated 4i workflow, we assessed the uniformity of antibody signal across 4i cycles. We performed 8 cycles of automated 4i, alternating between GFAP and ELAVL2 primary antibodies each cycle while the same secondary antibody cocktail was used across all cycles. The reproducibility of the GFAP and ELAVL2 expression pattern over 4i cycles was compared to the 1^st^ and 2^nd^ cycle images respectively. When compared to the 1^st^ cycle image, subsequent odd cycles faithfully reproduced the GFAP expression pattern while the signal was reduced to background in even cycles despite the presence of corresponding secondary antibody. Similarly, the ELAVL2 expression pattern was found to be consistent across even cycles, with a slight decrease in intensity in the 8th cycle, while the signal is reduced to background autofluorescence in odd cycles (Supplementary Figure 3). We demonstrated the ability to reproducibly stain and fully elute antibodies from tissue sections using our automated 4i protocol.

### Spectral unmixing enables 4 color imaging using the HiSeq 2500

We next sought to characterize the spectral properties of the HiSeq 2500 system in order to optimize 4 color automated imaging using PySeq2500. The HiSeq 2500 detects emission at the yellow-orange wavelengths 558 and 610nm from excitation with a green laser at 532 nm and emission at the far red wavelengths 687 and 740 nm from excitation with a red 660 nm laser. Commercially available fluorophore-conjugated secondary antibodies conventionally used for immunofluorescence typically have broad emission spectra that span multiple HiSeq detection channels, making simultaneous 4 color imaging a challenge. We identified fluorophores that are compatible with the HiSeq excitation source and emission filter (eg. Alexa Fluor 532, Alexa Fluor 594, Cy5, and Alexa Fluor 700) and tested crosstalk between them using a 4-channel imaging experiment. We used our automated 4i protocol to stain fresh frozen mouse spinal cord sections for astrocyte marker GFAP (AF594), neuronal marker ELAVL2 (AF700), nuclear envelope marker LMNB1 (AF532), and myelin marker MBP (Cy5) sequentially followed by staining for all four markers simultaneously in a fifth cycle. The fluorescence across emission channels for each singleplex cycle showed isolated signal from AF594 in the 610 nm channel and AF700 in the 740 nm channel. However, we observed significant spillover of AF532 from the 558 nm channel into the 610 nm channel and spillover of Cy5 signal from the 687 nm channel into the 740 nm channel (Figure 2A). For the multiplexed image generated in the fifth cycle, signal crosstalk was corrected using either linear unmixing or a blind source unmixing strategy based on PICASSO^21^ (Supplementary Figure 4).

**Figure 2.**
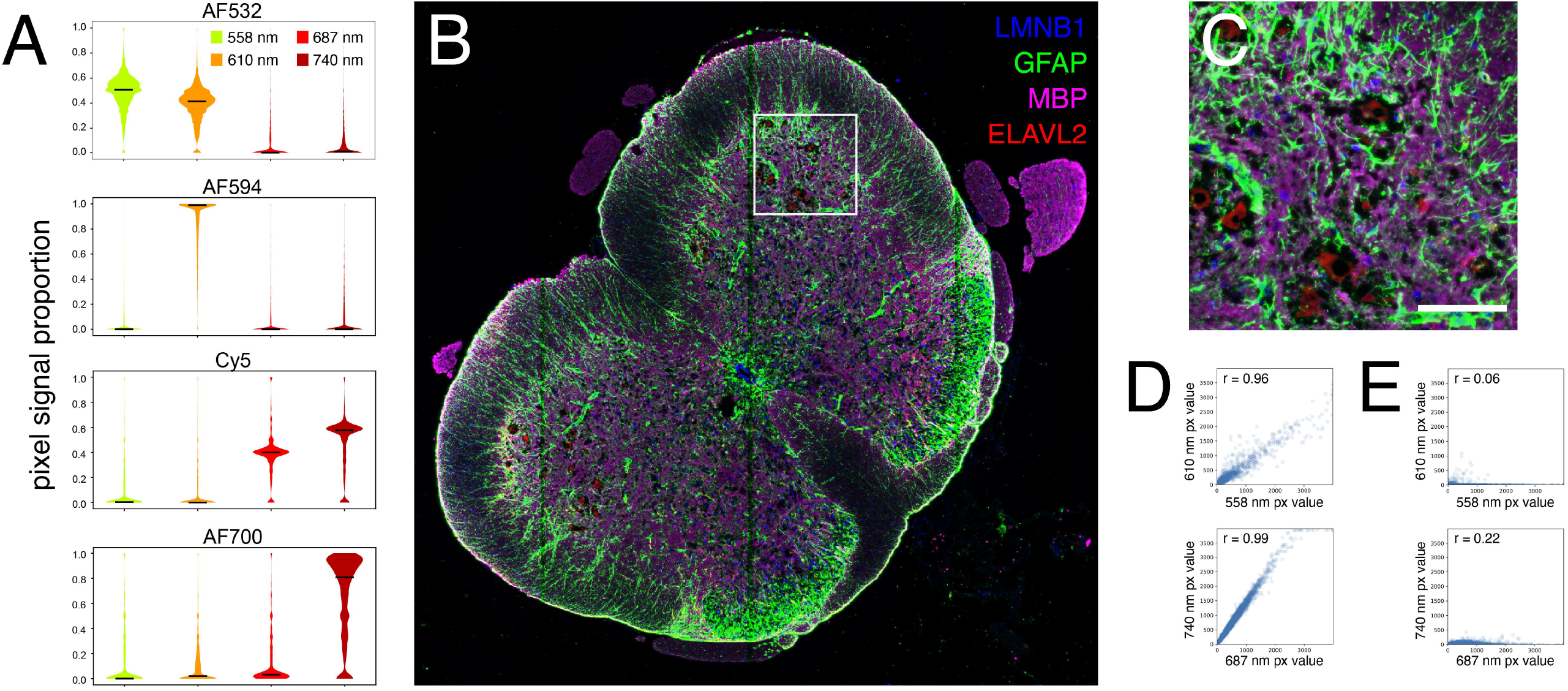
Simultaneous 4 channel imaging using PySeq2500. (A) Distribution of pixel signal intensity proportion across all detection channels from each fluorophore used for 4 channel imaging. (B) Mouse spinal cord stained simultaneously for LMNB1, GFAP, MBP, and ELAVL2 with fluorophore crosstalk removed using PICASSO unmixing strategy. (C) Inset of B. Scale bar represents 100 um. (D) Pearson correlation coefficient, r, of pixel intensities between corresponding detection channels prior to unmixing from singleplex staining of (top) LMNB1 with an AF532 conjugated secondary antibody and (bottom) MBP with a Cy5 conjugated secondary antibody. (E) Pearson correlation coefficient of pixel intensities between corresponding detection channels after linear unmixing of signal from singlex staining of (top) LMNB1 with an AF532 conjugated secondary antibody and (bottom) MBP with a Cy5 conjugated secondary antibody.

Blind source unmixing strategies, such as PICASSO, calculate the relative leakage between channels based on the multiplexed image alone. The PICASSO strategy iteratively minimizes the mutual information between image channels to calculate the optimal mixing matrix. We used a modified PICASSO strategy without iteration to unmix multiplexed images acquired in the fifth round of this experiment (Figure 2B-C, Supplementary Figure 5). In contrast to blind unmixing, linear unmixing requires reference images of individual stains for each detection channel to calculate crosstalk between channels. We used the signals of individual stains acquired in the first four cycles as reference images to calculate the relative leakage between channels from fluorophores that emitted in multiple channels. For example, the relative leakage of AF532 signal into the 610 nm detection channel was calculated by correlating pixel intensities from the 568 nm channel to the 610 nm channel image from the same cycle. The same procedure was used to measure the relative leakage of Cy5 signal from the 687 nm channel into the 740 nm channel (Figure 2D). The measured relative leakages were used to subtract the spillover fluorescence to unmix images with crosstalk (Figure 2E, Supplementary Figure 5). Linear unmixing and blind unmixing produced similar results. Therefore we used blind unmixing to correct images with spillover in subsequent experiments, eliminating the need to collect reference image data.

### Automated 4i generates replicable protein expression data

Automating the iterative chemistry and imaging steps required for 4i has the potential to greatly increase the throughput and reproducibility of multiplexed imaging experiments, enabling meaningful comparisons between biological conditions. To demonstrate the repeatability of protein expression measurements across technical replicates we performed automated 4i on serial mouse tissue sections processed i) in parallel on adjacent flow cells and ii) on separate runs. Adjacent spinal cord tissue sections from the SOD1 G93A mouse model of ALS^22^ and nontransgenic mice were sectioned across three flow cells that were processed over two automated 4i runs. To visualize cell type composition in spinal cord sections from the SOD1 G93A ALS mouse model after onset of motor symptoms as well as control animals, we stained for the following cell type markers across four cycles of 4i: astrocyte marker GFAP, neuronal markers ELAVL2, MAP2, and NFH, microglial marker IBA1, and white matter marker MBP (Supplemental Figure 6). The nuclear envelope marker LMNB1 was also included in every round (Supplemental Figure 6). Using our fully automated 4i approach, we generated hi-plex imaging data with subcellular resolution from 18 full tissue sections over 2 experimental runs that took 6 days each (Figure 3A).

**Figure 3.**
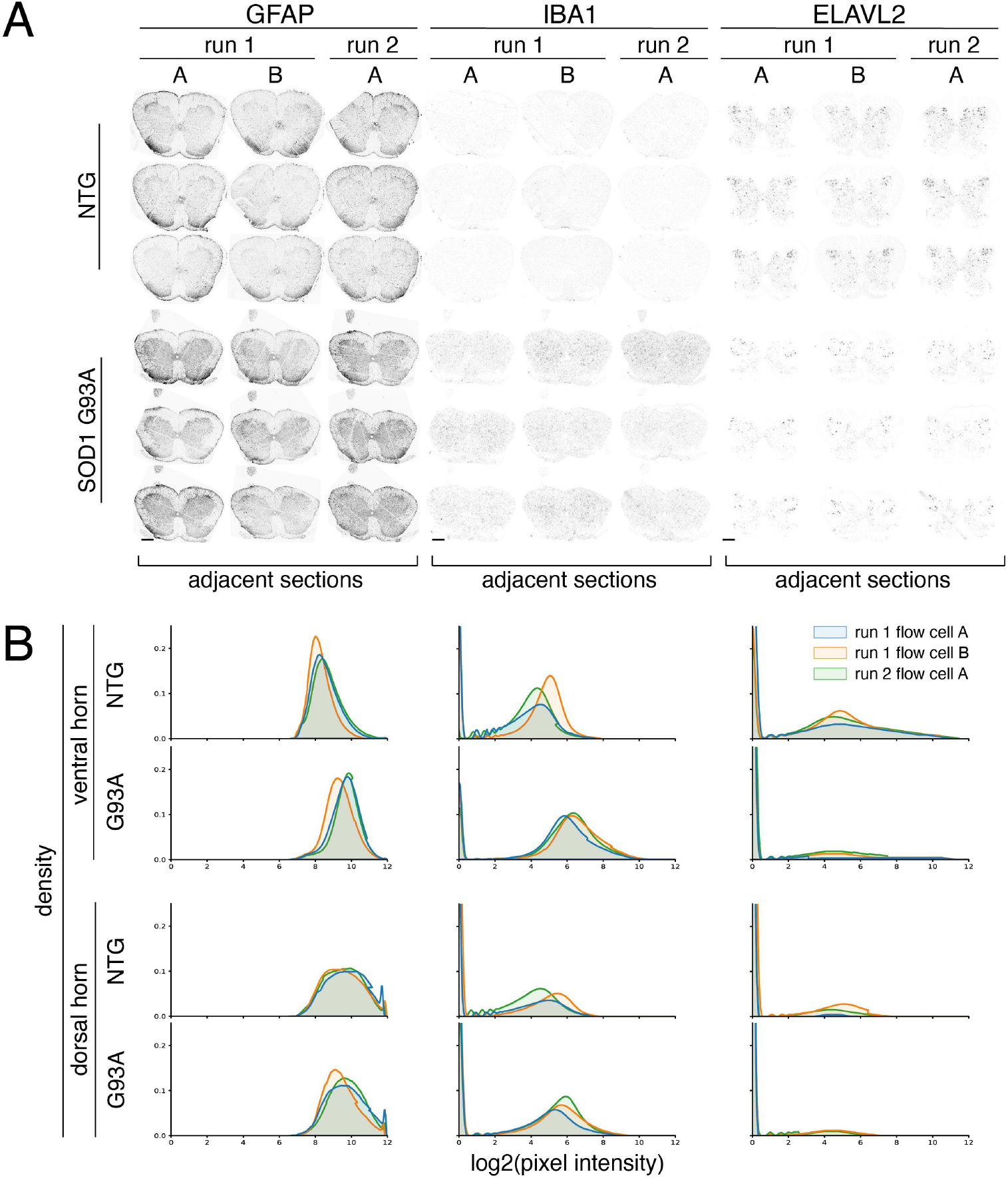
PySeq2500 automated 4i staining of adjacent spinal cord sections across flow cells and runs (A) Spinal cord sections from SOD1 G93A and nontransgenic animals stained for GFAP (astrocytes), IBA1 (microglia), and ELAVL2 (neurons) in two 4i cycles. Adjacent tissue sections from each animal were processed on flow cells run in parallel (run 1A and 1B) and across multiple experiments (run 1 and run 2). Scale bars represent 500 µm. (B) Histograms of signal intensity for each marker in manually annotated dorsal and ventral horn regions colored by experimental run.

PySeq2500 automation of the 4i protocol generated repeatable protein expression patterns on adjacent tissue sections across flow cells and runs (Figure 3A). Since loss of motor neurons from the ventral horn and reactive gliosis are known changes found in spinal cords from animals with the SOD1 G93A mutation after symptom onset^22,23^, we quantified the distribution of pixel intensities of ELAVL2, GFAP, and IBA1 in manually annotated ventral and dorsal horn regions of each tissue section. We observe similar pixel intensities for these stains between adjacent tissue sections processed in parallel and on different PySeq2500 runs (Figure 3B). Variability in pixel intensity within genotype is minimal while substantial differences in pixel intensities are found between genotypes. These results demonstrate the potential of automated reagent handling and imaging using PySeq2500 to generate repeatable data between technical replicates, enabling biologically meaningful comparisons between samples.

### Postmortem human spinal cord

To demonstrate the utility of PySeq2500 for automation of complex iterative reagent handling and imaging protocols over several days on large tissue sections, we also performed hi-plex immunostaining of fresh frozen postmortem tissue sections from ALS patients. The ability to multiplex unmodified, off the shelf antibodies as part of the 4i protocol allowed us to customize a panel of antibodies that label cell type markers and clinically validated pathognomonic protein inclusions found in spinal cords from patients with ALS^24–27^. We used these tools to jointly visualize neurons and neuronal processes (ELAVL3, MAP2, NFH), astrocytes (GFAP, ALDH1L1, AQP4), microglia (IBA1, CD68), myelin (MBP), vasculature (CD34) and pathological inclusions (pTDP-43, TDP-43, and p62) in postmortem spinal cords from patients with ALS (Figure 4). To enable visualization of 14 channel image data, we subjected the resulting image stacks to k-means clustering and assigned pixels in each cluster a color corresponding to its cluster label. This visualization approach reveals structure within 14 channel data that is not evident when viewing individual channels or when overlaying a subset of channels (Figure 4A). Using PySeq2500, we imaged cellular and pathological features at micron resolution across 2 tissue sections in each of 2 flowcells simultaneously. The combined tissue area in a single such experiment is over 350mm^2^. This data acquisition occurred over six cycles of fully automated staining, imaging, and elution. As such, our novel flow cell allowed reagent exchange and imaging while preserving tissue integrity over the 5 day experiment. Due to the robustness of the PySeq2500 system, we are able to register images acquired in sequential rounds of 4i to build a hi-plex view of tissue composition.

**Figure 4.**
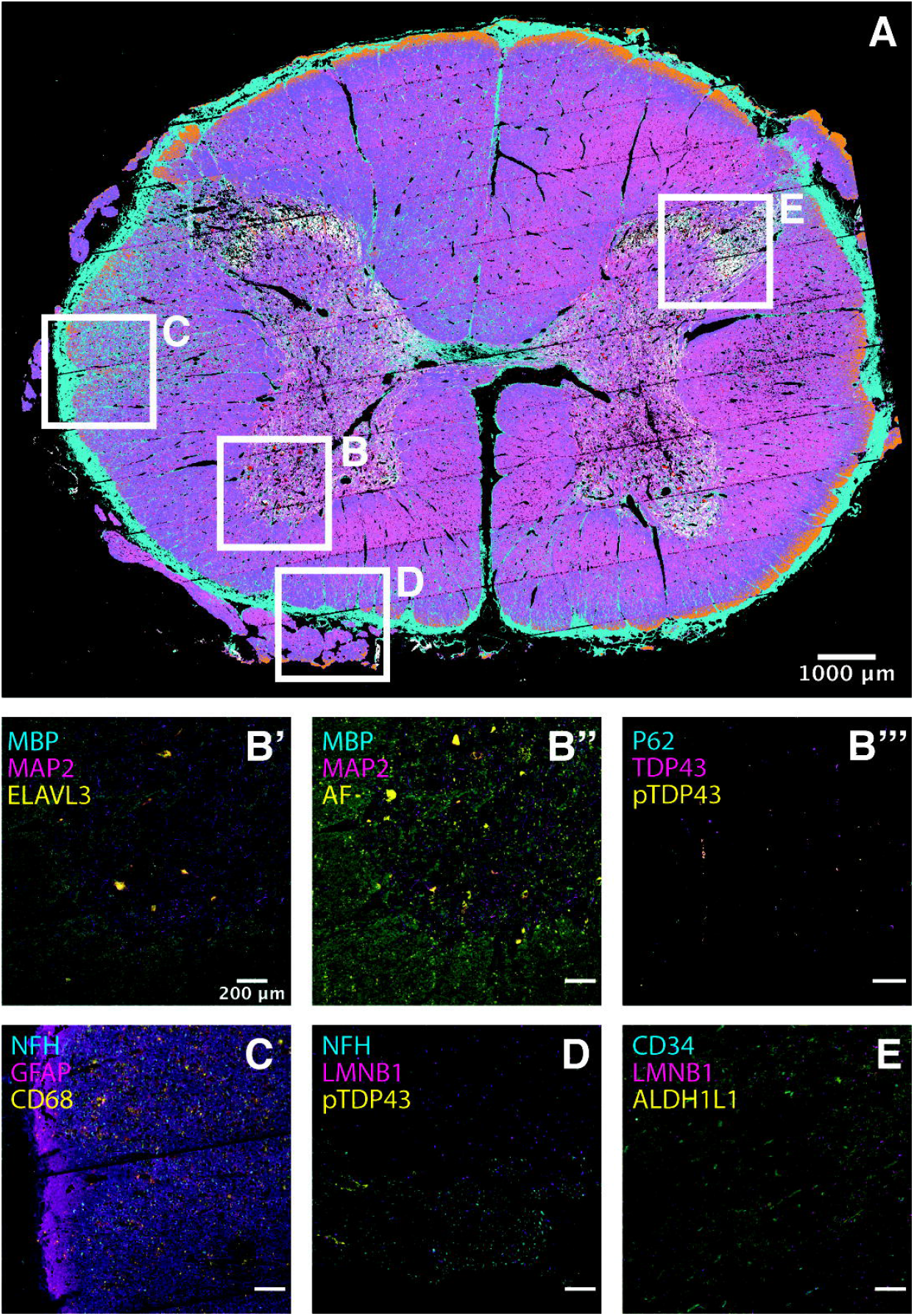
High-plex 4i tissue profiling of human postmortem spinal cord. Fresh frozen tissue sections from ALS patient spinal cords were subjected to 14-plex profiling over six 4i cycles. (A) K-means clustering of 4i data was used to visualize tissue domains defined by all 14 channels across the entire tissue section. Each of 7 clusters is assigned a unique color and inset locations are noted (B-E). (B’) Higher magnification image of anterior horn inset region B. Cell type marker MBP (cyan) is expressed by oligodendrocytes, MAP2 (magenta) is expressed in neurons and neuronal processes, and ELAVL3 (yellow) is expressed by neurons, including motor neurons. (B’’) Same as B’, but yellow is used to visualize tissue autofluorescence. (B’’’) Pathological markers P62 (cyan), TDP-43 (magenta), and hyperphosphorylated TDP43 (yellow) are visualized in the same FOV as in B’ and B’’. (C) Higher magnification image of lateral white matter inset region C. Cell type markers NFH expressed in neurons (cyan), GFAP expressed in astrocytes (magenta), and CD68 expressed in activated microglia (yellow) are visualized. (D) Higher magnification image of lateral white matter inset region D. Cell type markers NFH expressed in neurons (cyan), IBA1 expressed in microglia (yellow), and nuclear envelope marker LMNB1 (magenta), are visualized. (E) Higher magnification image of dorsal horn inset region E. Cell type markers CD34 expressed in endothelial cells (cyan), ALDH1L1 expressed in astrocytes (yellow), and nuclear envelope marker LMNB1 (magenta), are visualized.

## Discussion

We used automated 4i to demonstrate the capabilities of PySeq2500, an open source and customizable software toolkit and flow cell that enables unattended execution of demanding methods requiring precise control of many parameters simultaneously. Our automated 4i protocol generates repeatable protein expression maps that enable biologically relevant measurements of cell type composition and protein localization across conditions. We also demonstrate the application of automated 4i using PySeq2500 to generate high dimensional data of large postmortem tissue sections from patients with ALS. Our use of mouse and human spinal cord tissue sections also demonstrates the utility of our flow cell design for analysis of tissues with widely varying dimensions, compositions, and physical properties. Further, we demonstrate remarkable tissue stability for the extended period over which 4i experiments are conducted and through the accompanying fluid dynamics during the many reagent exchange steps involved. The robustness and repeatability of data generated using our automated 4i protocol demonstrates the potential of PySeq2500 to increase the throughput and replicability of imaging based -omics data generation.

Beyond immunofluorescence based applications, PySeq2500 has the potential to automate other protocols using iterative reagent exchange and imaging including *in situ* sequencing of transcriptomes and genomes. PySeq2500 could also be used to generate oligonucleotide arrays for solid phase capture based methods. Cho et al. suggest that their SeqScope technique could be scaled from the MiSeq flow cell to HiSeq 2500 flow cells, thereby increasing the surface area to 90 mm^2^ for solid phase capture based spatial transcriptomics^7^. Our PySeq2500 software and flow cell design makes this possible with a capture area that is approaching the size of a full slide (1875 mm^2^) and not limited by the lane geometry or availability of Illumina flow cells. Further, our flow cell was designed with accessible materials and can be readily customized for alternative applications by drilling holes to align with additional ports on the HiSeq stage and cutting the polyimide tape to create multiple chambers or chambers of different geometries. PySeq2500 is under active development and available on the Python Package Index as pyseq2500 and at https://github.com/nygctech/PySeq2500. We anticipate that our open source approach to developing the PySeq2500 toolkit will reduce the barrier to entry for additional labs to conduct spatially resolved and high throughput -omics scale experiments.

## Methods

### HiSeq 2500 modification

The stage was modified by removing the chamfered corner at the back left of each flow cell slot to allow common 25 × 75 mm glass slides to fit. To accommodate the two-hole flow cell design and increase the number of reagent lines for automated 4i, each syringe pump was modified by joining the inlets of barrels 1-4 and 5-8 to the stage outlets 4 and 5 respectively with a 5 port PEEK manifold (VICI, C5M1PK), ¼-28 flanged tube fittings with washers (VICI, CF-1A and CF-W1), and 1/16” OD PTFE tubing. Therefore, 2 mLs of liquid could be pulled from the central stage outlets with 1 pump stroke and stage outlets 1-3 and 6-8 were not in use. Additional reagent lines were connected to unused reagent valve ports using tubing and fittings from a discarded HiSeq 2500. Alternatively, 1/16 in. outer diameter PTFE tubing and 6-40 one-piece nuts/bushings (VICI, CNNF1PK) can be used. An approximately 1 cm diameter hole was drilled on the left side of the HiSeq 2500 to run the extra reagent lines to reagent reservoirs located outside of the HiSeq.

### Flow cell stage temperature

The HiSeq 2500 stage temperature of the A and B flow cell slots were set to a sequence of setpoints, held for 90 s at each setpoint, and measured by the HiSeq and an external thermocouple. The order of setpoints for the A flow cell was 25, 35, 45, 55, 25, 55, and 25° C. The order of setpoints for the B flow cell was 25, 35, 45, 55, 20, 55, and 20° C.

### Murine ALS model

B6SJL SOD1-G93A transgenic and nontransgenic control mice were obtained from Jackson Laboratories (Bar Harbor, ME), and maintained in full-barrier facilities at Columbia University Medical Center in accordance with ethical guidelines established and monitored by Columbia University Medical Center’s Institutional Animal Care and Use Committee. SOD1-G93A mice were monitored closely for onset of disease symptoms, including hindlimb weakness and weight loss.

### Spinal cord collection and sectioning

Mice were transcardially perfused with 1X Phosphate buffered saline (PBS) followed by spinal cord dissection. The L1-L3 and L3-L5 lumbar regions were isolated based on ventral root anatomy and embedded in Optimal Cutting Temperature (OCT, Fisher Healthcare, USA). The samples were then plunged into a bath of dry ice and pre-chilled ethanol until frozen and stored at - 80°C. Cryosections were cut at 10 μm thickness onto HiSeq compatible slides and stored at -80°C until use.

### Postmortem samples

Postmortem lumbar spinal cord samples from sporadic ALS patients were obtained from the Target ALS Multicenter Postmortem Core (http://www.targetals.org/). Cryosections were cut at 10 μm thickness onto HiSeq compatible slides and stored at -80°C until use.

### Tissue preparation

Mouse frozen sections were incubated on an Eppendorf Thermomixer with Smartblock adapter set to 37°C for 30 seconds. Sections were post-fixed with 10% formalin for 30 minutes at room temperature. Slides were washed six times with phosphate buffered saline. Sections were permeabilized with 0.5% Triton X-100 for 15 minutes at room temperature. Slides were washed six times with phosphate buffered saline. Frozen sections from postmortem ALS patients were incubated on an Eppendorf Thermomixer with Smartblock adapter set to 37°C for 1 minute followed by air drying at room temperature for 30 minutes. Sections were post-fixed with 10% formalin for 30 minutes at room temperature. Slides were washed six times with phosphate buffered saline. Sections were permeabilized with 0.1% Triton X-100 for 15 minutes at room temperature. Slides were washed six times with phosphate buffered saline.

### HiSeq flow cell assembly

Raw glass slides (25 × 75 mm) custom drilled with four 0.5 mm diameter holes that align with the ports on the stage of the HiSeq 2500 were obtained from Potomac Photonics according to the dimensions in Supplementary Figure 1A. After ultrasonic cleaning, slides were coated with 0.01% poly-l-lysine for 5 minutes and dried overnight at room temperature. Flow cell chambers were cut out of 4 mil (101.6 □m) thick polyimide tape with double sided adhesive backing (Bertech PPTDE-2) and adhered to the glass slide with the outer protective film intact. A Cricut Explore Air 2 cutting machine (Cricut 2004419) was used to cut the polyimide tape into the shape of the flow cell chamber using the vector file provided in Supplemental Materials. Samples were cryosectioned into the open flow cell and stored at -80°C until tissue processing. After tissue preparation as described above, the outer protective film was removed and a 25mm × 75mm 1.5 coverglass was adhered to the slide (Ibidi 10812).

### Elution optimization experiment

An automated PySeq2500 protocol was used to carry out the following steps: 1) fresh frozen mouse spinal cord sections were stained with ELAVL2 and GFAP and imaged, 2) the tissue was incubated with 4i elution buffer for 10 minutes 3) the elution buffer was exchanged for 4i imaging buffer and 4) the tissue was imaged. Steps 2-4 were repeated iteratively for 6 total elution cycles. The pixel intensity of each stain was measured after each elution cycle.

### Automated 4i protocol

A PySeq2500 configuration file was used to define reagent port and cycle assignments, pump settings, imaging parameters and ROIs, total number of cycles and the 4i recipe file. The 4i recipe file consisted of reagent exchange, hold, and imaging steps as described previously^4^. Mouse and human spinal cords were cryosectioned onto custom flow cells and processed as described above. After the flow cell was assembled, it was manually filled with phosphate buffered saline. The positions of tissue sections within the flow cell were manually measured and entered into the PySeq2500 configuration file. Blocking, imaging, and elution buffers were freshly prepared as described previously^4^. Antibody cocktails were prepared in conventional blocking buffer. Fluidics lines on the HiSeq 2500 were inserted into reagents as defined in the reagent port assignment section of the PySeq2500 configuration file. The HiSeq 2500 was initialized and all fluidics lines were primed with their corresponding reagents. Then experimental flow cells were locked onto the HiSeq 2500 stage and held in place with a vacuum. The PySeq2500 experiment was initiated and the software automatically stepped through reagent exchange, hold, and imaging steps as defined in the 4i recipe, applying imaging settings as defined in the configuration file. Briefly, each 4i cycle consisted of a one hour incubation with 4i blocking buffer, two hour incubations with primary and secondary antibody cocktails, imaging in 4i imaging buffer, and either four or five ten minute incubations with 4i elution buffer for mouse and human tissue sections respectively. Full once and partial once autofocus routines were used for the mouse and human sections respectively. Human tissue sections were imaged before the first round of staining to assess autofluorescent features. Antibodies used in each round for each experiment are reported in Supplemental Table 1.

### Antibodies

Detailed description of antibodies used in each cycle of each experiment are provided in Supplemental Table 1. Elution series and alternating 4i experiments were conducted using GFAP (Abcam, ab4674; Novus, NBP1-05198) and ELAVL2 (Atlas, HPA063001) antibodies. Simultaneous 4 color imaging was conducted using GFAP (Abcam, ab4674), ELAVL2 (Atlas, HPA063001), LMNB1 (Sigma, AMAB91251), and MBP (Abcam, ab209328) antibodies. Mouse spinal cord 4i data was generated over four cycles using LMNB1 (Sigma, AMAB91251), ELAVL2 (Atlas, HPA063001), GFAP (Abcam, ab4674), IBA1 (Abcam, ab178847), MAP2 (Abcam, ab5392), NFH (Abcam, ab4680), and MBP (Abcam, ab209328) antibodies. Data for PVALB (Swant, PV27a) and PDGFRa (Cell Signaling, 3174S) are not shown. Human spinal cord 4i data was generated over six cycles using CD34 (R&D Systems, AF7227), LMNB1 (Sigma, AMAB91251), CD68 (DAKO, M0814), TDP-43 (Proteintech, 10782-2-AP), MAP2 (Abcam, ab5392), ELAVL3 (Invitrogen, A-21271), pTDP-43 (Proteintech, 22309-1-AP), NFH (Abcam, ab4680), IBA1 (Abcam, ab178847), GFAP (Abcam, ab4647), MBP (Atlas, AMAb91062), ALDH1L1 (Atlas, HPA050139), and P62 (Abcam, ab56416) antibodies. Data for TMEM119 (Atlas, AMAb91528) and AQP4 (Atlas, HPA014784) antibodies) are not shown.

### Autofocus

The first step of the general procedure for the full, partial, full once and partial once autofocus routines is to obtain an out-of-focus image of the ROI. Then, the out-of-focus image is analyzed to find, filter and rank FOVs of interest based on distance and contrast. Objective stacks at top ranked FOVs are obtained by continuously imaging frames of the FOV as the objective moves away from the stage. The sharpness of each frame is estimated by computing the file size of the JPEG-compressed frame, which increases in relation to sharp image features. A curve is fitted to the JPEG-compressed file size as a function of objective position, and the objective position corresponding to the maximal file size is saved. A number of focal objective positions at multiple FOVs are calculated and the median value is used as the optimal objective position for the ROI.

### Raw image processing

PySeq2500 saves imaging tiles as 16 bit TIFFs with 12 bit pixel depth. Each of the 4 TDI line scanning CCD cameras consist of 8 groups of 256 pixels and each group of pixels, ***g***, registered a different average dark pixel value, ***d_px***_***g***,***camera***_. A dark pixel offset was calculated for each pixel group as the mean of ***d_px***_***g***,***camera***_ across the camera minus ***d_px***_***g***,***camera***_. Pixel group intensity values were adjusted by the group dark pixel offset to correct the background across an image. Shifts in corresponding pixels across channels due to chromatic aberration were found by imaging tetraspeck beads and manually measuring the pixel translation. Images were rigidly registered according to the measured shifts so that pixels across channels had a 1 to 1 correspondence. Images of human tissue section stains were additionally rigidly registered over cycles to the tissue autofluorescence image acquired before the first 4i cycle. The relative shift of cycles was measured using phase cross correlation with only the central 2.3 mm^2^ FOV of the ROI from the 610 nm channel to reduce computation time. The measured shifts were used to translate the images and align autofluorescence features in the tissue section.

### Unmixing of 4 color images

Pixel signal intensity proportion from singleplex stains were calculated first by selecting only corresponding pixels across channels above the background from a primary channel, where the primary detection channels for AF532, AF594, Cy5, and AF700 were the 558, 610, 687, and 740 nm channels respectively. Then the signal intensity of a pixel from a specific channel, ***px***_**i**,**channel**_, was divided by the sum of pixel intensities across all channels, ∑_□ □ □ □ □ □ □_ ***px***_**i**,**channel**_.

We used a modified PICASSO algorithm to estimate the relative leakage of signal, ***x***, from a fluorophore in one image (***D***_***i***_) to another image from an adjacent channel (***D***_***j***_). First we background subtracted images. The mode of pixel intensities plus 1 standard deviation was used as a global background pixel value for images of mouse sections. Autofluorescence images prior to any staining were used to subtract background from human sections. Then, the mutual information, ***I***, between ***D***_***j***_ and ***D***_***j***_***-xD***_***i***_ was computed at a range of ***x*** between 0 and 2. The optimal ***x*** was found from the minimal point of a curve fit to ***I***(***x***) and the unmixed image was calculated as ***D***_***j***_-***xD***_***i***_.

The relative leakage of signal by linear unmixing, ***x***_***l***_, was measured by taking reference images of a single fluorophore across channels. The reference images were background subtracted and then ***x***_***l***_ was estimated as the slope of a line fit to corresponding pixel intensities of ***D***_***i***_ and ***D***_***j***_ and weighted by ***D***_***i***_ intensity. The linearly unmixed image was calculated as ***D***_***j***_-***x***_***l***_***D***_***i***_.

Pearson correlation coefficients were calculated from 10000 random pixels above background in primary channels and corresponding pixels in adjacent channels.

### Histogram adjustments

Raw 12bit images were rescaled to fit 16bit encoding. For clarity of presentation, histograms were subjected to linear or gamma adjustment. When used, such adjustments were applied uniformly for each antibody within experiments.

### Histogram plotting

Mouse spinal cord 4i data was processed as described previously, registered across 4i cycles, and subjected to maximum intensity projection of 10 z-planes. Ventral and dorsal horns were manually annotated on the basis of MAP2 and ELAVL2 staining using Napari^28^. Histograms of pixel intensity were generated using 10^6^ randomly selected pixels per annotated ventral and dorsal horn areas per sample. Pixel intensity measurements were pooled between nontransgenic and SOD1 G93A technical replicates on each flow cell to allow comparisons between flow cells and genotypes. Density plots represent log2(pixel intensity). For clarity of representation of non-zero pixel intensities, background pixel intensities extend beyond the limits of the y-axis.

### K-Means Pixel clustering

14 channel 4i images were down sampled to 25% raw resolution to smooth any pixel scale registration defects using scikit-image’s rescale module^29^. Images were then subjected to K-means clustering using scikit-learn’s ‘cluster.kmeans’ module (n_clusters=7)^30^.

## Code Availability

PySeq2500 and is available at https://github.com/nygctech/PySeq2500.

## Acknowledgements

We would like to acknowledge the New York Genome Center for providing the retired HiSeq 2500 sequencing machines used in this work. We would also like to acknowledge Urs Gaudenz and the ReSeq Hackteria team for providing detailed information about the HiSeq 2000^17^. Work in H.P.’s lab is supported by the National Institutes of Health (NS117583, NS116350, NS118183, NS118570, HG011014, AG066831), CZI, the ALS Association, the Tow Foundation, and Target ALS.

## Author Contributions

KP wrote the PySeq2500 software and modified the HiSeq 2500, JP adapted the 4i protocol for tissue sections and executed automated 4i experiments, KP and JP designed the PySeq2500 flow cell, MC performed mouse husbandry and spinal cord isolation, KP, JP, and SM performed data analysis, and KP, JP, SM, WS, PS and HP conceived of experiments and wrote the manuscript. KP and JP contributed equally to this work, and have the right to list their names first on their CV.

## Competing Interests

The authors declare no competing interests.

## Data Availability

The datasets generated during and/or analysed during the current study are available from the corresponding author on reasonable request.

## Notes

https://github.com/nygctech/PySeq2500

### Competing Interest Statement

The authors have declared no competing interest.

