## Supplementary Materials for "PySeq2500: An open source toolkit for repurposing HiSeq 2500 sequencing systems as versatile fluidics and imaging platforms"

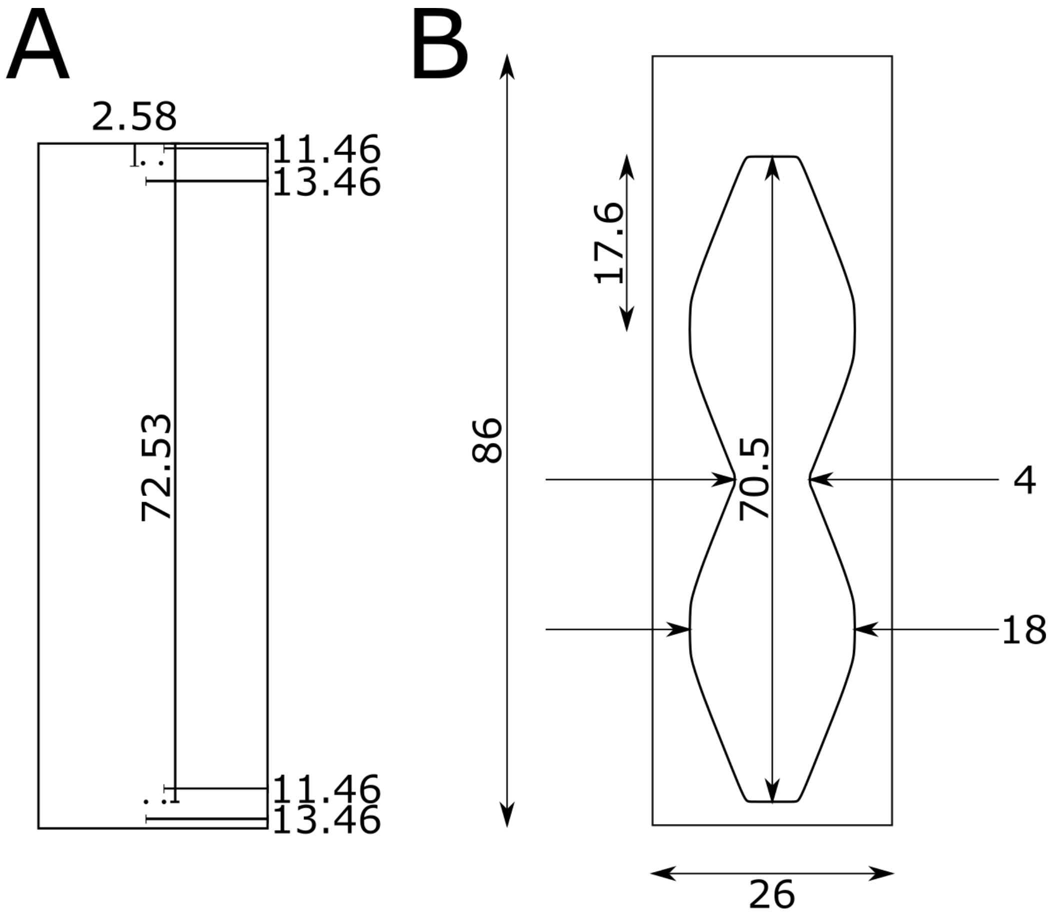


**Supplementary Figure 1** Design of a PySeq2500 flow cell compatible with the HiSeq2500 stage and integrated fluidics (A) Positions of 0.5 mm diameter through holes on a 25 x 75 mm glass slide which align with the 2 hole stage inlets and outlets 4 & 5 on the HiSeq 2500 stage. (B) PySeq2500 flow cell chamber dimensions. Flow cell chambers are manufactured using a Cricut cutting machine from 4 mil thick polyimide tape with double sided adhesive. Excess polyimide tape is trimmed to the edge of the slide before flow cell assembly. (all dimensions in mm)

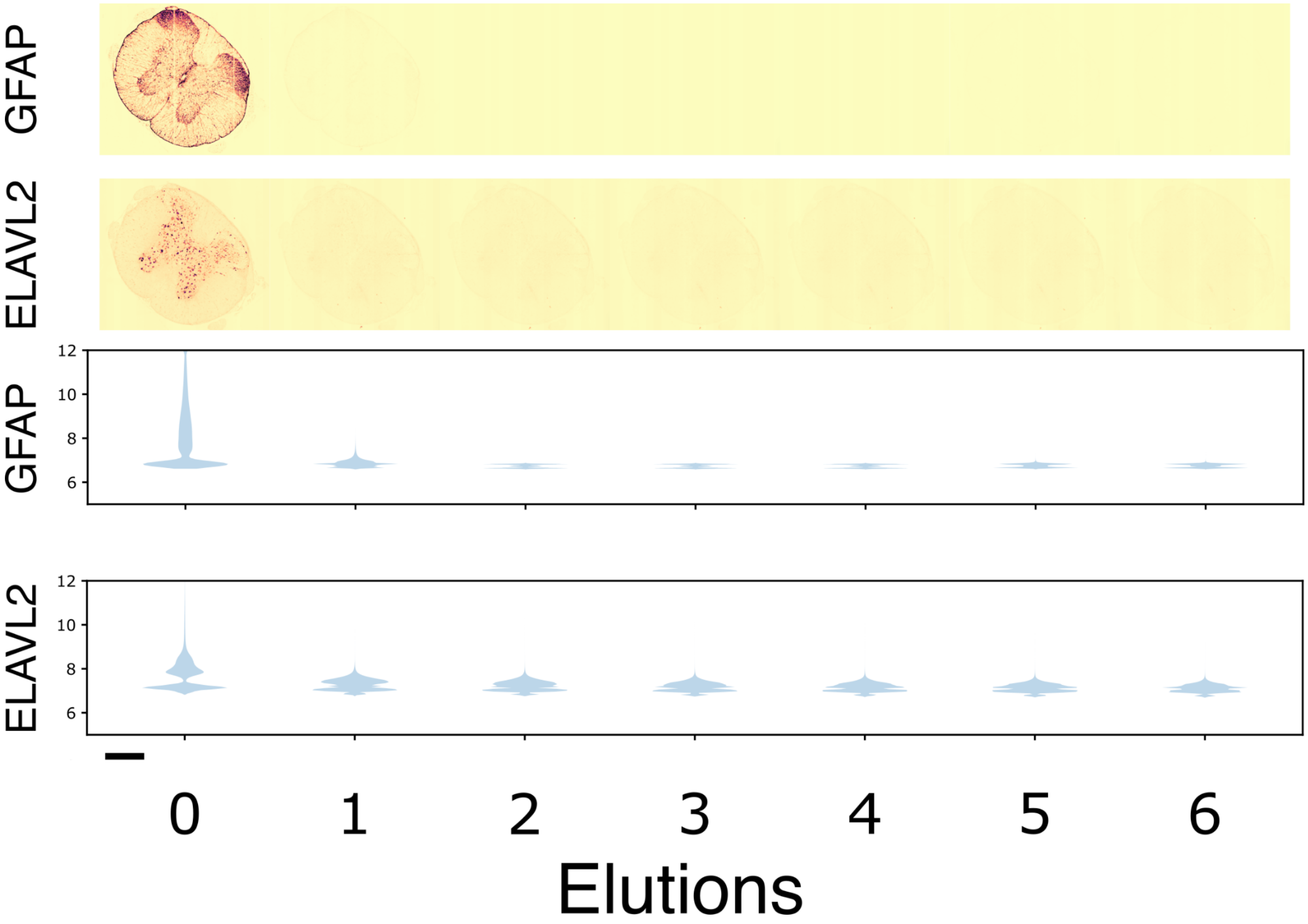


**Supplementary Figure 2** Automated elution series for optimizing 4i protocol for use on fresh frozen tissue sections. (*top*) A mouse spinal cord section was stained for GFAP (687 nm channel) and ELAVL2 (610 nm channel) and imaged after sequential rounds of elution. (*bottom*) Histograms of $10^6$ random pixels from each round of elution. Y axis represents log2(pixel intensity).

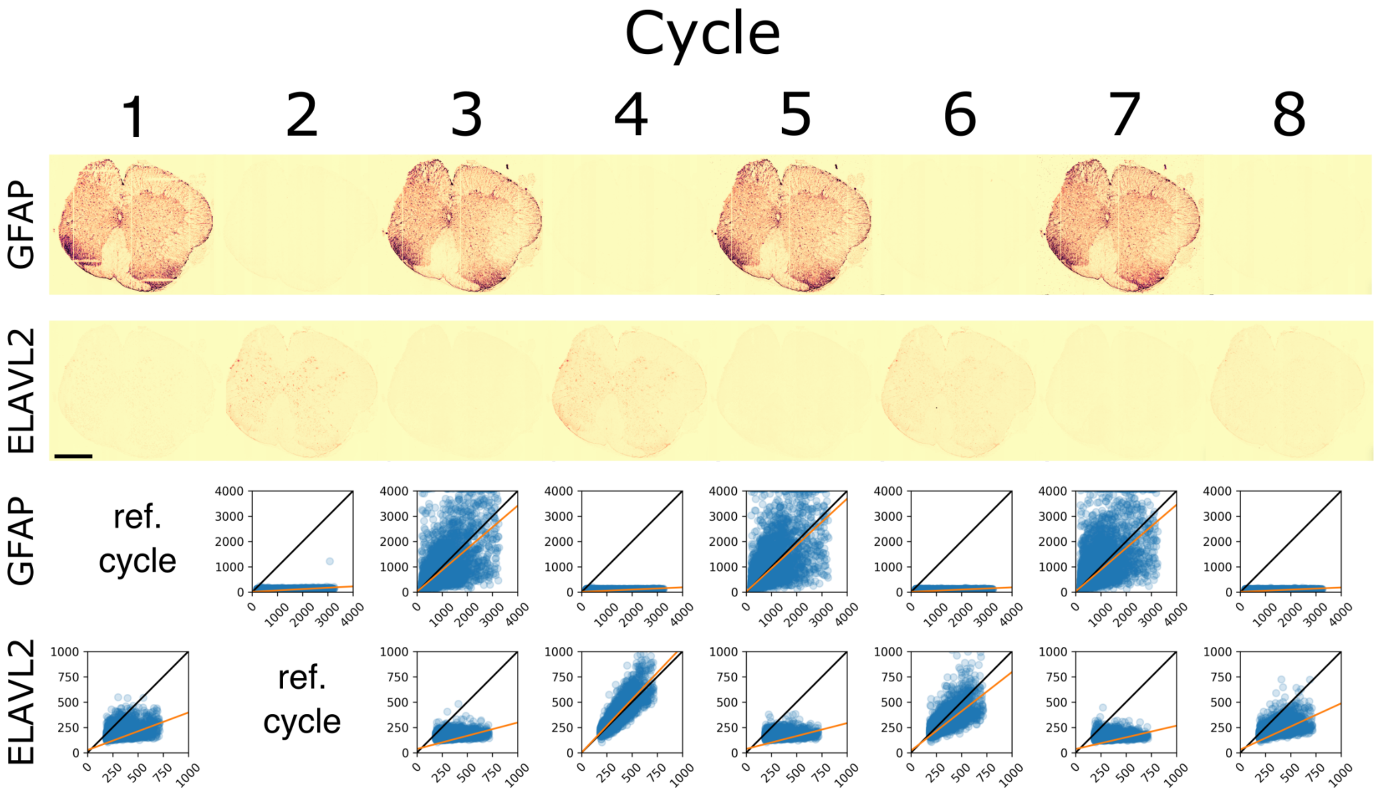


**Supplementary Figure 3** Mouse spinal cord section was immunostained with Chicken anti-GFAP in odd cycles, Rabbit anti-ELAVL2 in even cycles using our PySeq2500 automated 4i protocol. The same secondary antibody cocktail, Cy5 donkey anti-Chicken and AF594 Donkey anti-Rabbit was used in each of the 8 cycles. (*top*) Images of GFAP (687 nm channel) and ELAVL2 (610 nm channel) from each cycle of staining (scale = 500 µm). (*bottom*) Randomly subsampled ($10^5$ pixels) pixel intensities from the reference cycle vs all other cycles. The first cycle of staining with each primary antibody, cycles 1 and 2 for GFAP (687 nm channel) and ELAVL2 (610 nm channel) respectively, was used as the reference cycle. All pixels fit to a line weighted by pixel intensity for each cycle is shown as an orange line and y = x is shown as a black line.

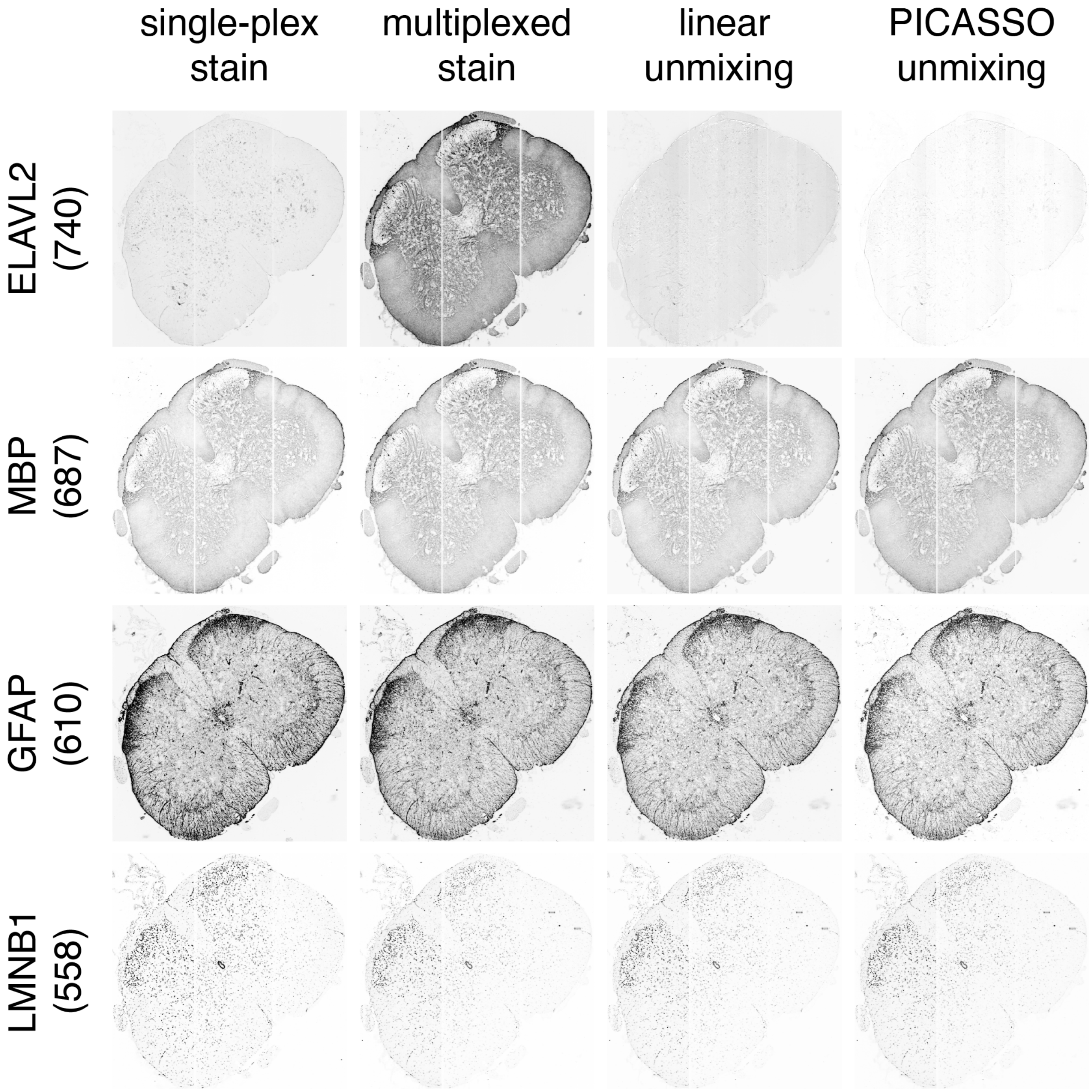


**Supplementary Figure 4** Mouse spinal cord section stained in singleplex cycles (cycles 1-4) and in the mixed multiplexed cycle (cycle 5). Individual stains from the multiplexed cycle (cycle 5) calculated using linear unmixing and PICASSO unmixing strategies.

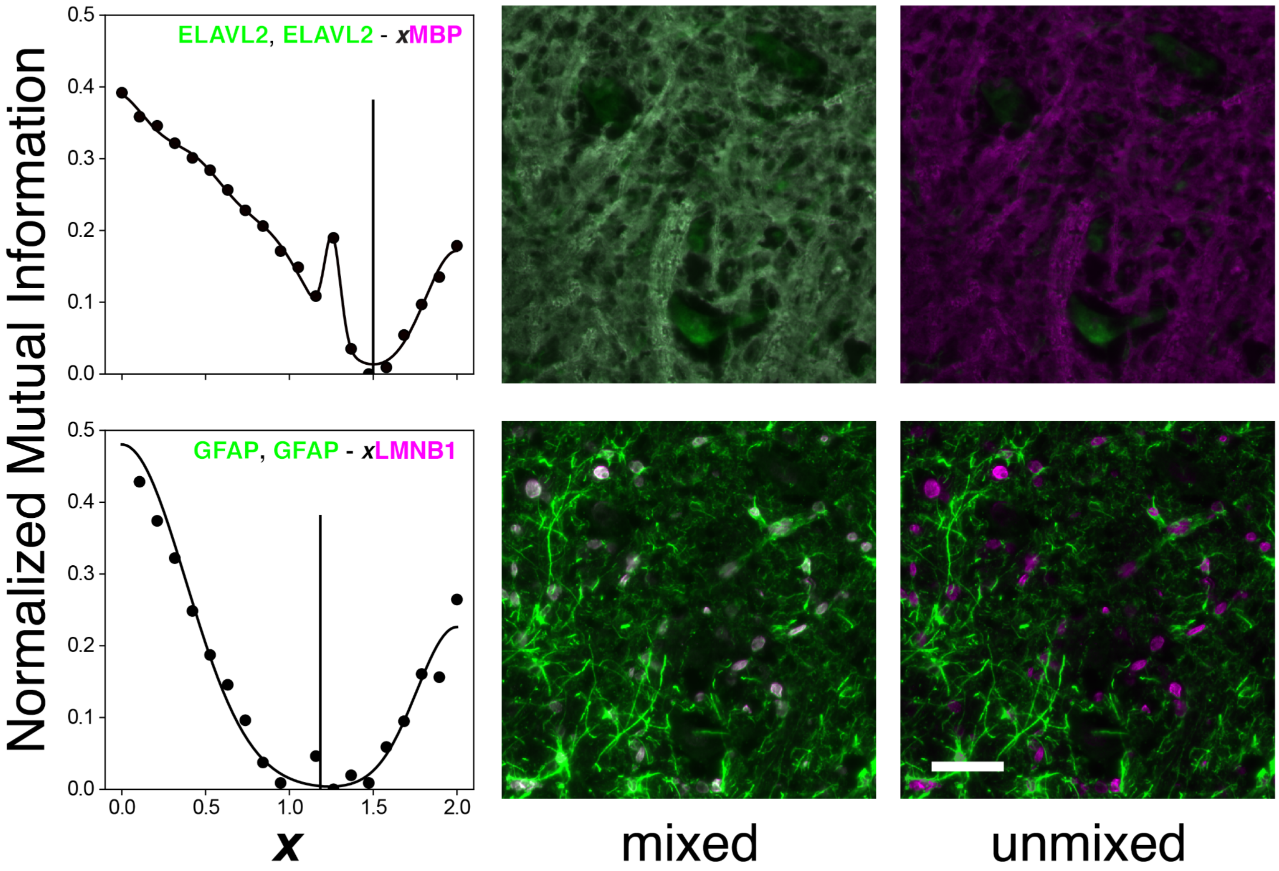


**Supplementary Figure 5** PICASSO unmixing by minimization of mutual information. Mutual information, ***I***, was calculated at a range of relative leakage, ***x***, from (*top*) MBP into ELAVL2 and (*bottom*) LMNB1 into GFAP. The optimal ***x*** was estimated as the minimal point of a curve fit to ***I***(***x***) and used to unmix the spillover signal. Scale bar represents 100 um.

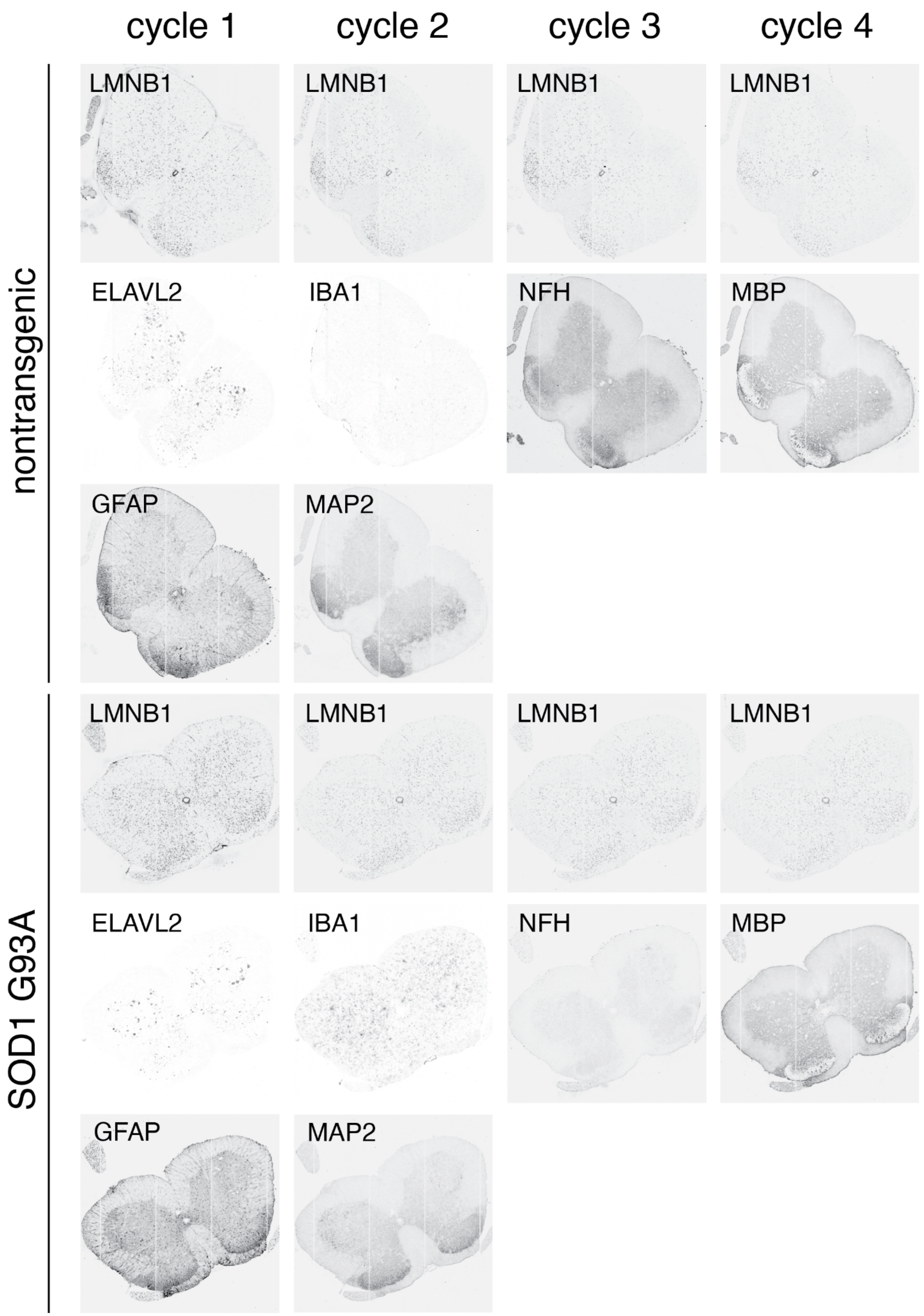


**Supplementary Figure 6** Fresh frozen spinal cord sections from nontransgenic and SOD1 G93A mutant mice were subjected to four cycles of 4i to visualize six cell type markers, with LMNB1 in each cycle.

| FIGURE | TISSUE | 4i CYCLE | PRIMARY | VENDOR | CATALOG # | CLONE | HOST | DILUTION | LOT # | SECONDARY | VENDOR | CATALOG # | DILUTION | LOT# |
|---|---|---|---|---|---|---|---|---|---|---|---|---|---|---|
| Supplemental figure 2 | WT p70, lumbar L1-L3, fresh frozen | Cycle 1 | GFAP | Abcam | ab4674 | polyclonal | Chicken | 1:500 | GR3368950-1 | Cy5 Donkey Anti-Chicken | Jackson | 703-175-155 | 1:500 | 130611 |
| | | | ELAVL2 | Atlas | HPA063001 | polyclonal | Rabbit | 1:100 | 000015755 | Alexa Fluor 594 Donkey Anti-Rabbit | Jackson | 711-585-152 | 1:500 | 140419 |
| Supplemental figure 3 | WT p70, lumbar L1-L3, fresh frozen | Odd cycles | GFAP | Novus | NBP1-05198 | polyclonal | Chicken | 1:500 | 7529-18 | Cy5 Donkey Anti-Chicken | Jackson | 703-175-155 | 1:500 | 130611 |
| | | | | | | | | | | Alexa Fluor 594 Donkey Anti-Rabbit | Jackson | 711-585-152 | 1:500 | 140419 |
| | | Even cycles | ELAVL2 | Atlas | HPA063001 | polyclonal | Rabbit | 1:100 | 000015755 | Cy5 Donkey Anti-Chicken | Jackson | 703-175-155 | 1:500 | 130611 |
| | | | | | | | | | | Alexa Fluor 594 Donkey Anti-Rabbit | Jackson | 711-585-152 | 1:500 | 140419 |
| Figure 2, Supplemental figures 4-5 | WT p30, lumbar L1-L3, fresh frozen | Cycle 1 | GFAP | Abcam | ab4674 | polyclonal | Chicken | 1:500 | GR3368950-2 | Alexa Fluor 594 Goat Anti-Chicken | Invitrogen | A11042 | 1:500 | 673785 |
| | | Cycle 2 | ELAVL2 | Atlas | HPA063001 | polyclonal | Rabbit | 1:100 | 000015755 | Alexa Fluor 700 Goat Anti-Rabbit | Invitrogen | A21038 | 1:500 | 2129003 |
| | | Cycle 3 | LMNB1 | Sigma | AMAB91251 | CL3929 | Mouse | 1:100 | MAB-03076 | Alexa Fluor 532 Goat Anti-Mouse | Invitrogen | A11002 | 1:500 | 2160043 |
| | | Cycle 4 | MBP | Abcam | ab209328 | IGX3421 | Human | 1:1000 | GR3215004-10 | Cy5 Donkey Anti-Human | Jackson | 709-175-149 | 1:500 | 129880 |
| | | Cycle 5 | LMNB1 | Sigma | AMAB91251 | CL3929 | Mouse | 1:100 | MAB-03076 | Alexa Fluor 532 Goat Anti-Mouse | Invitrogen | A11002 | 1:500 | 2160043 |
| | | | GFAP | Abcam | ab4674 | polyclonal | Chicken | 1:500 | GR3368950-2 | Alexa Fluor 594 Goat Anti-Chicken | Invitrogen | A11042 | 1:500 | 673785 |
| | | | MBP | Abcam | ab209328 | IGX3421 | Human | 1:1000 | GR3215004-10 | Cy5 Donkey Anti-Human | Jackson | 709-175-149 | 1:500 | 129880 |
| | | | ELAVL2 | Atlas | HPA063001 | polyclonal | Rabbit | 1:100 | 000015755 | Alexa Fluor 700 Goat Anti-Rabbit | Invitrogen | A21038 | 1:500 | 2129003 |
| Figure 3, Supplemental figure 6 | M387, NTG p100 male; M388 SOD1 G93A p100 male | Cycle 1 | LMNB1 | Sigma | AMAB91251 | CL3929 | Mouse | 1:200 | MAB-03076 | Alexa Fluor 532 Goat Anti-Mouse | Invitrogen | A11002 | 1:500 | 2160043 |
| | | | ELAVL2 | Atlas | HPA063001 | polyclonal | Rabbit | 1:100 | 000015755 | Alexa Fluor 594 Donkey Anti-Rabbit | Jackson | 711-585-152 | 1:1000 | 151658 |
| | | | GFAP | Abcam | ab4674 | polyclonal | Chicken | 1:500 | GR3368950-3 | Cy5 Donkey Anti-Chicken | Jackson | 703-175-155 | 1:1000 | 152944 |
| | | Cycle 2 | LMNB1 | Sigma | AMAB91251 | CL3929 | Mouse | 1:200 | MAB-03076 | Alexa Fluor 532 Goat Anti-Mouse | Invitrogen | A11002 | 1:500 | 2160043 |
| | | | IBA1 | Abcam | ab178847 | EPR16589 | Rabbit | 1:250 | GR3229566-15 | Alexa Fluor 594 Donkey Anti-Rabbit | Jackson | 711-585-152 | 1:1000 | 151658 |
| | | | MAP2 | Abcam | ab5392 | polyclonal | Chicken | 1:500 | GR3386708-1 | Cy5 Donkey Anti-Chicken | Jackson | 703-175-155 | 1:1000 | 152944 |
| | | Cycle 3 | LMNB1 | Sigma | AMAB91251 | CL3929 | Mouse | 1:200 | MAB-03076 | Alexa Fluor 532 Goat Anti-Mouse | Invitrogen | A11002 | 1:500 | 2160043 |
| | | | PVALB | Swant | PV27a | polyclonal | Rabbit | 1:1000 | NA | Alexa Fluor 594 Donkey Anti-Rabbit | Jackson | 711-585-152 | 1:1000 | 151658 |
| | | | NfH | Abcam | ab4680 | polyclonal | Chicken | 1:250 | GR3359372-1 | Cy5 Donkey Anti-Chicken | Jackson | 703-175-155 | 1:1000 | 152944 |
| | | Cycle 4 | LMNB1 | Sigma | AMAB91251 | CL3929 | Mouse | 1:200 | MAB-03076 | Alexa Fluor 532 Goat Anti-Mouse | Invitrogen | A11002 | 1:500 | 2160043 |
| | | | PDGFRa | Cell Signaling | 3174S | D1E1E | Rabbit | 1:500 | 8 | Alexa Fluor 594 Donkey Anti-Rabbit | Jackson | 711-585-152 | 1:1000 | 151658 |
| | | | MBP | Abcam | ab209328 | IGX3421 | Human | 1:1000 | GR3215004-10 | Cy5 Donkey Anti-Human | Jackson | 709-175-149 | 1:500 | 129880 |
| Figure 4 | sporadic ALS patients | Cycle 1 | CD34 | R&D Systems | AF7227 | | Sheep | 1:50 | CFZV0121051 | Cy5 Donkey Anti-Sheep | Jackson | 713-175-147 | 1:1000 | 152945 |
| | | | LMNB1 | Sigma | AMAB91251 | CL3929 | Mouse | 1:100 | MAB-03502 | Alexa Fluor 594 Donkey Anti-Mouse | Jackson | 715-585-150 | 1:1000 | 153991 |
| | | Cycle 2 | CD68 | DAKO | M0814 | KP1 | Mouse | 1:100 | 41258320 | Alexa Fluor 532 Goat Anti-Mouse | Invitrogen | A11002 | 1:500 | 2160043 |
| | | | TDP-43 | Proteintech | 10782-2-AP | polyclonal | Rabbit | 1:250 | 00065465 | Alexa Fluor 594 Donkey Anti-Rabbit | Jackson | 711-585-152 | 1:1000 | 151658 |
| | | | MAP2 | Abcam | ab5392 | polyclonal | Chicken | 1:500 | GR3386708-1 | Cy5 Donkey Anti-Chicken | Jackson | 703-175-155 | 1:1000 | 152944 |
| | | Cycle 3 | ELAVL3 | Invitrogen | A-21271 | 16A11 | Mouse | 1:250 | 2105721 | Alexa Fluor 532 Goat Anti-Mouse | Invitrogen | A11002 | 1:500 | 2160043 |
| | | | pTDP-43 | Proteintech | 22309-1-AP | polyclonal | Rabbit | 1:250 | 00058641 | Alexa Fluor 594 Donkey Anti-Rabbit | Jackson | 711-585-152 | 1:1000 | 151658 |
| | | | NFH | Abcam | ab4680 | polyclonal | Chicken | 1:250 | GR3359372-4 | Cy5 Donkey Anti-Chicken | Jackson | 703-175-155 | 1:1000 | 152944 |
| | | Cycle 4 | TMEM119 | Atlas | AMAb91528 | CL8714 | Mouse | 1:250 | MAB-03453 | Alexa Fluor 594 Donkey Anti-Mouse | Jackson | 715-585-150 | 1:1000 | 153991 |
| | | | IBA1 | Abcam | ab178847 | EPR16589 | Rabbit | 1:250 | GR3229566-20 | Alexa Fluor 700 Goat Anti-Rabbit | Invitrogen | A21038 | 1:500 | 2129003 |
| | | | GFAP | Abcam | ab4674 | polyclonal | Chicken | 1:500 | GR3393187-1 | Cy5 Donkey Anti-Chicken | Jackson | 703-175-155 | 1:1000 | 152944 |
| | | Cycle 5 | MBP | Atlas | AMAb91062 | CL2819 | Mouse | 1:500 | MAB02927 | Alexa Fluor 594 Donkey Anti-Mouse | Jackson | 715-585-150 | 1:1000 | 153991 |
| | | | ALDH1L1 | Atlas | HPA050139 | polyclonal | Rabbit | 1:1000 | 000016054 | Alexa Fluor 647 Donkey Anti-Rabbit | Jackson | 711-605-152 | 1:1000 | 154880 |
| | | Cycle 6 | P62 | Abcam | ab56416 | polyclonal | Mouse | 1:100 | GR3294261-1 | Alexa Fluor 594 Donkey Anti-Mouse | Jackson | 715-585-150 | 1:1000 | 153991 |
| | | | AQP4 | Atlas | HPA014784 | polyclonal | Rabbit | 1:1000 | 000014721 | Alexa Fluor 647 Donkey Anti-Rabbit | Jackson | 711-605-152 | 1:1000 | 154880 |

**Table 1** Primary and secondary antibodies used in automated 4i experiments.
